# Genetic Regulation of Circular RNA Expression in Human Aortic Smooth Muscle Cells and Vascular Traits

**DOI:** 10.1101/2022.01.26.477946

**Authors:** Redouane Aherrahrou, Dillon Lue, Mete Civelek

## Abstract

**Background:** Circular RNAs (circRNAs) are a class of non-coding RNAs that have cell-type specific expression and are relevant in cardiovascular disease. Aortic smooth muscle cells (SMCs) play a crucial role in cardiovascular disease by differentiating from a quiescent to proliferative phenotype. The role of circRNAs in SMCs and their relevance to cardiovascular disease is largely unexplored.

**Results:** In this study, we employ a systems genetics approach to identify circRNA transcripts at a genome wide level and their relevance in cardiovascular traits. We quantified circRNA expression across 151 quiescent and proliferative human aortic SMCs from multiethnic donors. We identified 1,589 expressed circRNAs. Between quiescent and proliferative SMCs, we identified 173 circRNAs which were differentially expressed. To characterize the genetic regulation of circRNA expression, we associated the genotypes of 6.3 million single nucleotide polymorphisms (SNPs) with circRNA abundance and found 96 circRNAs which were associated with genetic loci. Three SNPs were associated with circRNA expression in proliferative SMCs but not in quiescent SMCs. We identified 6 SNPs which had distinct association directions with circRNA isoforms from the same gene. Lastly, to identify the relevance of circRNAs in cardiovascular disease, we overlapped genetic loci associated with circRNA expression with vascular disease related GWAS loci. We identified 7 blood pressure, 1 myocardial infarction, and 3 coronary artery disease loci which were associated with a circRNA transcript (*circZKSCAN1, circFOXK2, circANKRD36, circLARP4, circCEP85L, circGTF3C2, circPDS5A, circSLC4A7, and chr17:42610108*|*42659552*) but not mRNA transcript.

**Conclusions:** Overall, our results provide mechanistic insight into the regulation of circRNA expression and the genetic basis of cardiovascular disease.

## INTRODUCTION

Smooth muscle cells (SMCs) are the major cellular component of the blood vessel wall. In physiological conditions, SMCs exist primarily in a contractile phenotype which allows them to maintain vascular homeostasis through vasodilation or contraction and proper vessel wall elasticity by secreting extracellular proteins.^1^ During pathophysiological conditions, SMCs differentiate from a contractile (quiescent) to a synthetic (proliferative) phenotype which reduces their ability to contract, alters the composition of secreted extracellular proteins, and increases their capacity for proliferation, migration, and calcification.^1^ This phenotypic switching results in vascular remodeling and therefore plays a role in hypertension, aortic aneurysms, coronary artery disease (CAD), strokes, and heart attacks.^2^

CircRNAs are an abundant, tissue-specific class of non-coding RNAs formed through a covalent linkage between the 5’ and 3’ end of an RNA molecule. CircRNAs have numerous functions including: acting as microRNA and RNA binding protein sponges, modulating the expression of linear transcripts, and being translated themselves.^3,4,5,6^ Because of their lack of ends, circRNAs are protected from exonucleases and have a longer life span than their linear counterparts.^7^ *CircANRIL*^8^, *circACTA2*^9^, *circLRP6*^10^, *circSFMBT2*^11^, and *circMAP3K5*^12^ are involved in SMC migration, proliferation, and phenotypic modulation. Further, the most significantly associated genomic locus with increased risk for CAD, 9p21, has been shown to regulate the expression of circular isoform of the long noncoding RNA, ANRIL in atherosclerotic plaques.^8^ However, a comprehensive study linking genetic variation to smooth muscle circRNA expression is missing.

Genome wide association studies (GWAS) have linked genetic loci with many cardiovascular traits including CAD, stroke, and high blood pressure.^13^ The mechanism of individual loci is largely unknown. Majority of the significantly associated loci map to non-coding regions of the genome pointing to their effect on regulating gene expression. Large scale studies such as the Genotype Tissue Expression (GTEx) project map GWAS loci to mRNA expression levels in what is known as expression quantitative trait loci (eQTL) analyses. The integration of GWAS and eQTL studies enables the prediction of causal genes affecting complex phenotypic traits.

In a recent study, we had found that 37% of CAD GWAS loci are associated with mRNA gene expression levels in aortic SMCs.^14^ In this study, we extended the traditional eQTL study to identify variants associated with circular RNA (circRNA) expression in aortic SMCs in what we term a circQTL (**Figure 1**). We cultured and performed RNA-sequencing on 151 SMCs in a quiescent and proliferative SMCs representing the contractile and synthetic phenotype respectively. We identified 1,589 circRNAs many of which were SMC phenotype specific. From these 1,589 circRNAs, we identified 139 genetically regulated circRNAs, three of which were phenotype specific. Using colocalization analyses, we identified 6 blood pressure, 1 myocardial infarction and 3 CAD GWAS loci which had association with circRNA expression but not mRNA expression. This study reveals that the genetic risk for cardiovascular diseases may be partially regulated by circRNAs.

**Figure 1:**
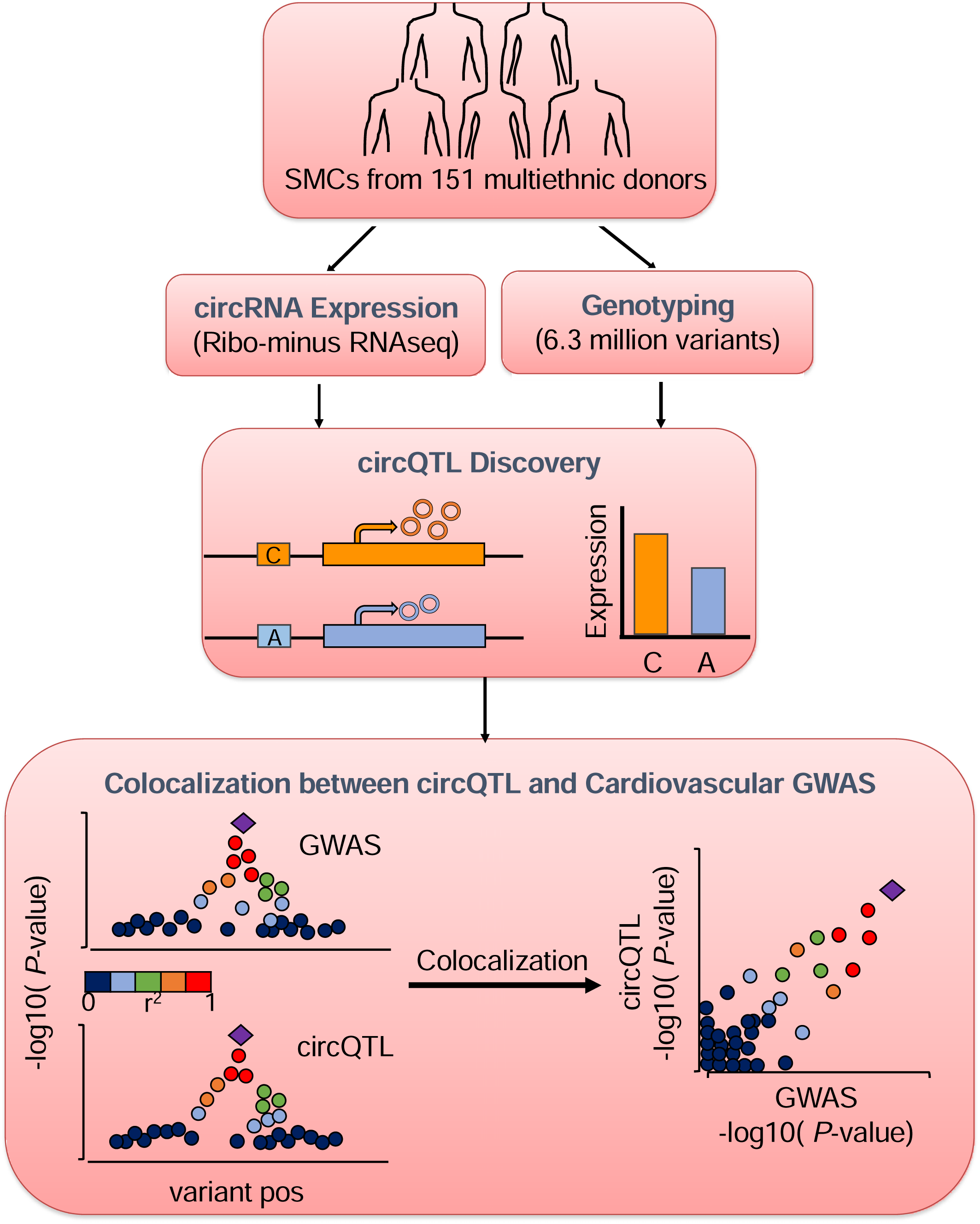
Study Design and Overview of Analysis. We quantified circRNA expression using RNA-seq of quiescent and proliferative human aortic smooth muscle cells (SMCs) from 151 multi-ethnic donors. We associated circRNA expression with ∼6.3 million variants to perform circRNA quantitative trait loci (circQTL) mapping. Subsequently, we performed colocalization between circQTLs and vascular GWAS traits.

## RESULTS

### Landscape of Circular RNA

We performed RNA sequencing on aortic SMCs derived from 151 ethnically diverse healthy heart transplant donors (118 male and 33 female). After quantification and quality control, we obtained 145 and 139 samples cultured in the presence (proliferative) or absence (quiescent) of FBS, respectively.^14^ We used CIRI^15^ to de novo identify and quantify circRNA backsplicing sites (**Figure 2A**). Although we assumed that each backsplicing site represents a unique circRNA transcript, a backsplicing site could represent multiple circRNA isoforms with differences in internal splicing. We identified 1,214 and 1,566 expressed circRNAs (junction counts ≥3 and the circular ratio ≥0.05 in at least 20% of the samples) in the quiescent and proliferative SMCs, respectively. From the two phenotypes combined, we identified 1,589 unique circRNAs (**Figure 2B**). A particular circRNA has a corresponding parent gene if the circRNA’s backsplicing site overlaps an annotated exon or intron of a gene. Out of the expressed circRNA transcripts, about 95% had a protein coding parent gene and only 2.1% had no parent gene (**Table 1**). Most circRNAs were exonic (92.5%) followed by intronic (5.3%) and intergenic (2.2%) (**Supplementary Figure 1**).

**Figure 2:**
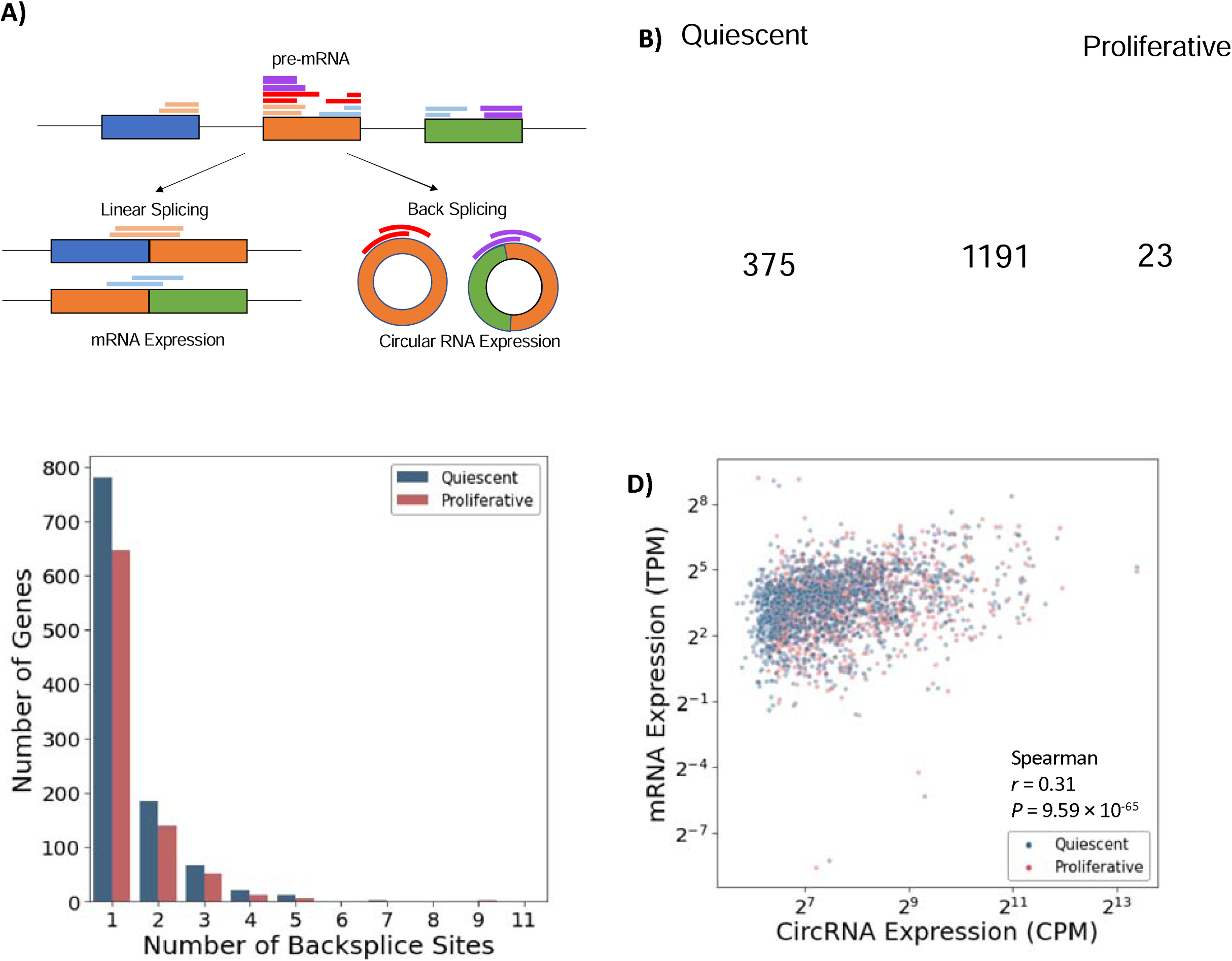
Circular RNA profile in human aortic smooth muscle cells. **A)** mRNAs are made through linear splicing where a splice acceptor appears downstream of the splice donor. In contrast, circRNAs are made through backsplicing where the acceptor site appears upstream of the donor site creating a circular molecule. circRNAs are detected by mapping RNA-seq reads (rectangles above exon) that overlap a backsplice site. The red and purple reads support two different circRNA transcripts respectively from the same gene. **B)** Venn diagram comparing the number of expressed circRNAs in quiescent versus proliferative SMCs. CircRNAs were considered expressed if the junction counts ≥3 and the ratio of mRNA reads to backsplice reads was ≥0.05 in at least 20% of samples. **C)** Histogram of number of transcripts (represented by the number of backsplice sites) per gene. **D)** Correlation between circRNA expression and mRNA expression of the corresponding parent gene.

**Table 1.** Number of circRNAs and circQTLs identified.

|  |  | **circRNA Expression** |  |
| --- | --- | --- | --- |
| **Phenotype** | **Parent Gene Type** | **Number of circRNA tested** | **Number of circRNA with circQTL** |
| **Quiescent** | protein coding | 1490 (95.1%) | 86 (90.0%) |
|  | lncRNA | 25 (1.6%) | 3 (3.1%) |
|  | pseudogene | 12 (0.8%) | 0 (0.0%) |
|  | Multiple Parent Gene | 6 (0.4%) | 2 (0.21%) |
|  | No Parent Gene | 33 (2.1%) | 5 (5.2%) |
|  | Total | 1566 | 96 |
| **Proliferative** | protein coding | 1151 (94.8%) | 81 (84.4%) |
|  | lncRNA | 19 (1.6%) | 4 (4.2%) |
|  | pseudogene | 10 (0.8%) | 4 (4.2%) |
|  | Multiple Parent Gene | 5 (0.4%) | 2 (2.1%) |
|  | No Parent Gene | 29 (2.4%) | 5 (5.2%) |
|  | Total | 1214 | 96 |

Additionally, we found that >70% of parent genes, had a single backsplicing site (**Figure 2C**). In our RNA seq data, we identified 18,964 expressed mRNAs, 1083 of which had a circRNA suggesting that 6% of SMC genes could be circularized. There was a statistically significant but low correlation (*r* = 0.31) between circRNA expression and its parent gene expression (**Figure 2D**).

### Differential circRNA Expression

Of the 1,589 expressed circRNAs, 173 were differentially expressed between the quiescent and proliferative phenotypes, with 62 circRNAs having a higher expression in quiescent SMCs and 111 circRNAs having a higher expression in proliferative SMCs. (**Figure 3A, Supplementary Table 1**). Using topGO, we tested the enrichment in gene ontology terms for the 139 parent genes of differentially expressed circRNAs against a background of SMC genes.^16^ The parent genes of the differentially expressed circRNAs were enriched in processes related to SMC contraction including: regulation of sodium ion transmembrane transport, regulation of heart contraction, and actin cytoskeleton organization (**Figure 3B**).

**Figure 3:**
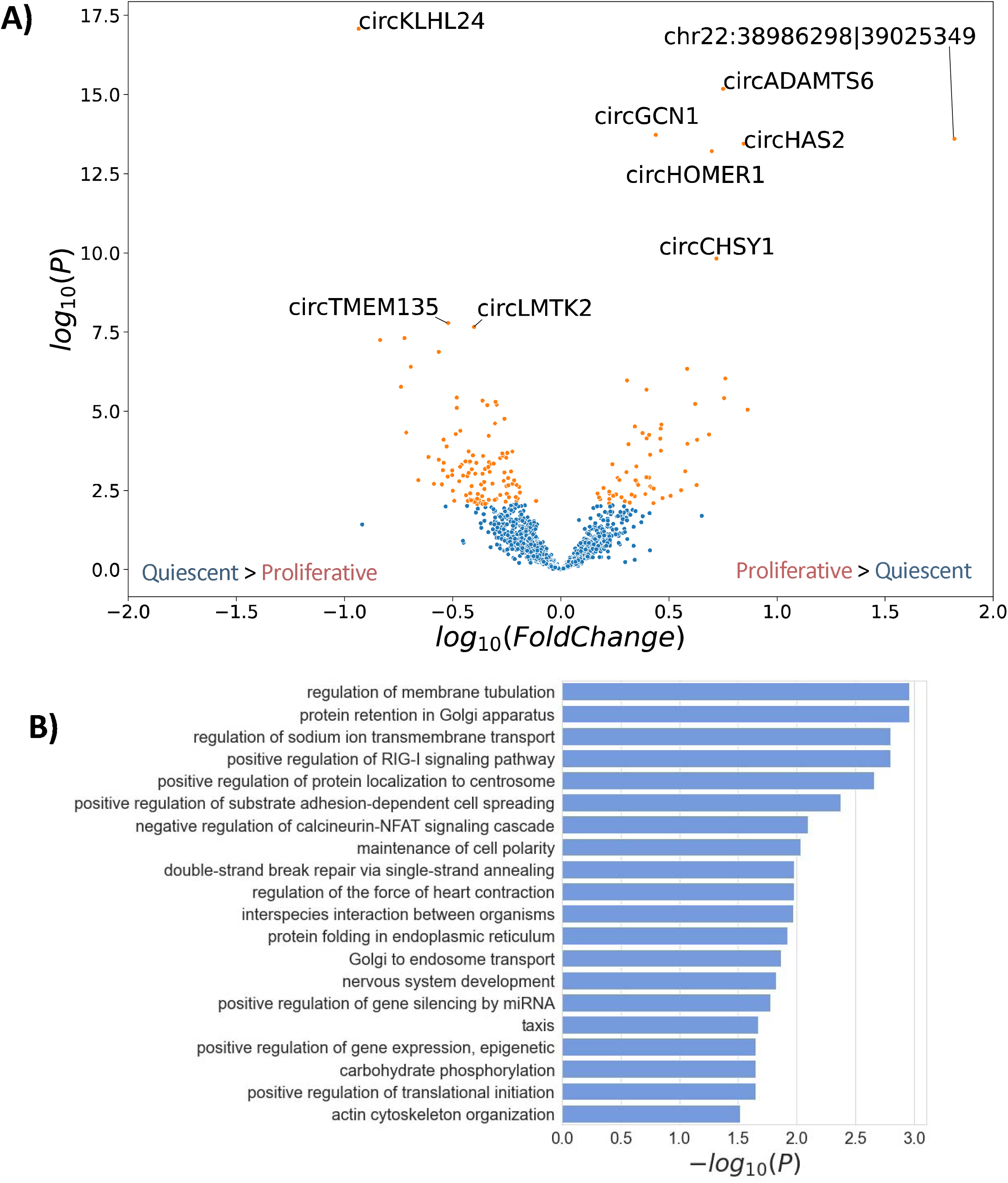
CircRNA differential expression between quiescent and proliferative SMCs. **A)** Volcano plot representing differentially expressed circRNAs. CircRNAs enriched in quiescent SMCs are further to the left and circRNAs enriched in proliferative SMCs are further to the right. The orange points represent the 173 circRNAs that passed significance *q*-value < 0.05. The top 10 most significant circRNAs are labeled. **B)** Gene ontology enrichment analysis of the parent genes of differentially expressed circRNAs. All terms with *P* < 0.05 are shown.

### Quantitative Trait Loci Mapping

We next tested the association between circRNA expression with single nucleotide polymorphisms (SNPs) at least 500 kb away from the backsplicing site in each SMC phenotype separately using tensorQTL. 8 and 4 PEER covariates maximized the number of circQTLs identified.^17^ Of the 1,589 expressed circRNAs tested, we identified 96 circQTLs in both SMC phenotypes separately and 53 circQTLs which were common to both SMC phenotypes (**Figure 4A, Supplementary Table 2**). For each circQTL, we termed the most significant or lead SNP in the locus as eSNP. After conditioning on the eSNP of each circQTL, we identified one secondary circQTL in quiescent SMCs and two in proliferative SMCs. 86% of the circRNAs with circQTL had a parent protein-coding gene. Using topGO, we tested the enrichment in gene ontology terms for the 139 circQTL parent genes against a background of SMC genes.^16^ We identified enrichment in various pathways associated with SMC biology including Rho protein signal transduction, sodium ion transmembrane transporter activity, and cAMP-mediated signaling (**Figure 4B**).

**Figure 4:**
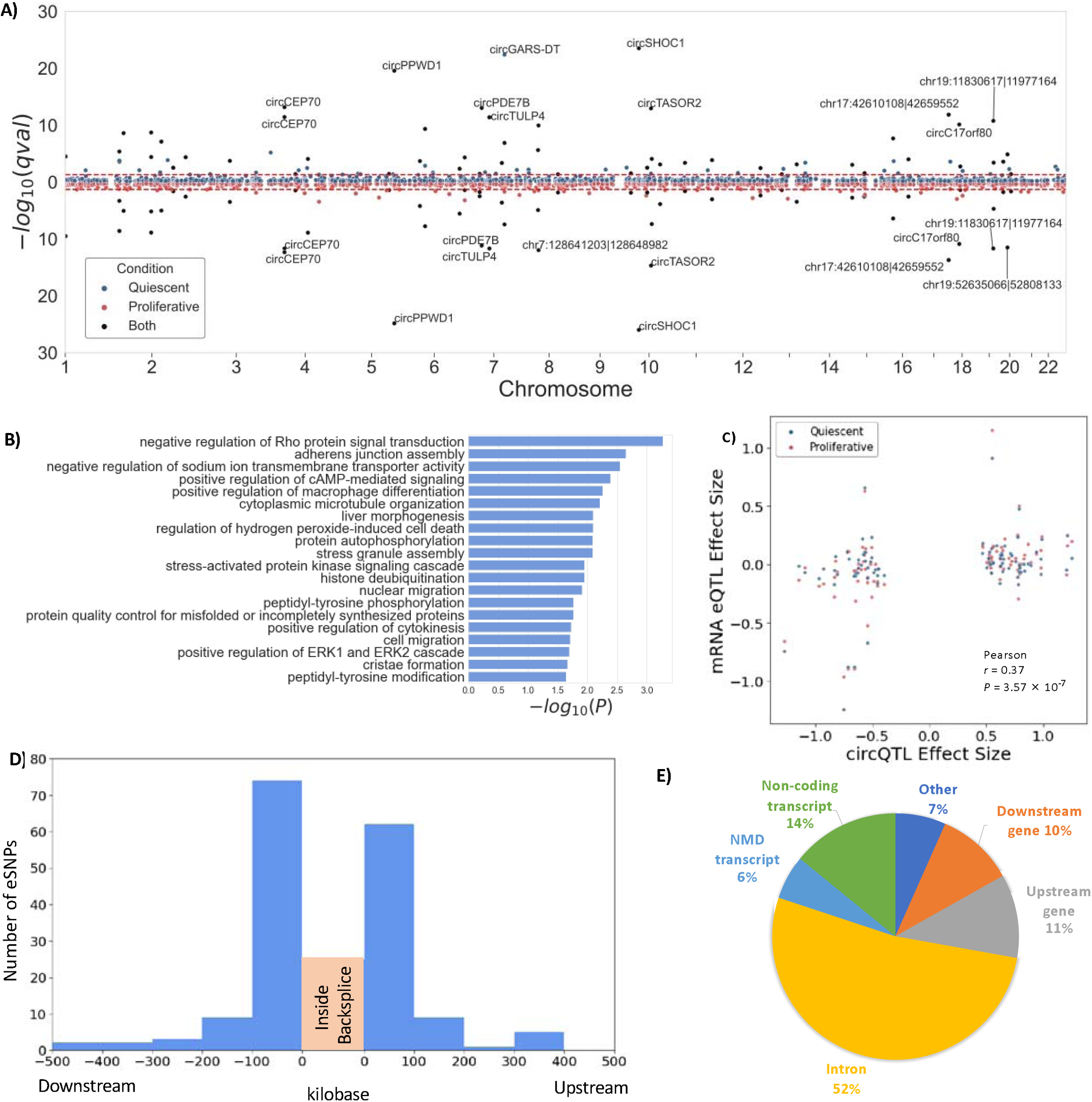
Genetic regulation of circRNA expression. **A)** Miami plot showing the most significant association between a circRNA and a variant within a locus. Top half represents quiescent circQTLs while the bottom half represents proliferative circQTLs. The red line indicates the significance level threshold at *q*-value=0.05. All circRNAs with *q*-value less than 1× 10^−48^ are labeled. **B)** Gene ontology enrichment analysis of the parent genes of circQTLs in either proliferative or quiescent SMCs. All terms with *P* < 0.05 are shown. **C)** Pearson correlation between SNP effect size on circRNA expression and SNP effect size on parent gene mRNA expression. **D)** Histogram of distances of the eSNPs from backsplice site. **E)** Annotated effects of eSNPs using Variant Effect Predictor.

Next, we tested whether the effect size of circQTL SNPs on circRNA expression had similar effect size on mRNA expression of its parent gene. We calculated the correlation of the eSNP effect size on circRNA expression and eSNP effect size on parent gene expression. We determined low but significant correlation (*r* = 0.37, P= 3.57 × 10^−7^) (**Figure 4C**). Although simple, this analysis did not consider the many SNPs in high linkage disequilibrium within a genetic locus. To formally test whether a genetic locus is associated with circRNA expression but not mRNA expression of the parent gene, we performed colocalization for each significant circQTL locus with the eQTL locus using COLOC.^18^ Across the 179 circQTL loci with parent genes in quiescent and proliferative SMCs, we identified 26 loci which were associated with mRNA expression of their parent gene. From each SMC phenotype, we highlighted an example of a colocalized and not colocalized locus (**Supplementary Figure 2)**. To annotate circQTL SNPs, we determined the distribution of eSNP distance from the backsplicing site and found that 82% of eSNPs were within 100 kb of the circRNA backsplicing site (**Figure 4D**). Additionally, we found that 53% eSNPs were within introns and 14.1% were in non-coding transcripts (**Figure 4E**).

### Phenotype-specific circQTLs

We previously showed that phenotypic state of SMCs affect the genetic regulation of mRNA expression;^14^ therefore, we tested for circQTLs that were specific to either proliferative or quiescent SMCs. For each eSNP, we correlated the effect size of the eSNP on circRNA expression in the proliferative and quiescent SMCs. We found low correlation (*r* = 0.36, P=7.01 × 10^−48^) indicating phenotype-specific genetic regulation (**Figure 5A**). Having identified low correlation, we formally tested differences in eSNP effect size between SMC phenotypes. We identified three circQTLs (*circSTRN3, circCHD9*, and *circRIPK1*) that had a significant association with a SNP in one phenotype but not the other phenotype (**Figure 5B,C,D**). Statistical test results for each circRNA are in **Supplementary Table 3**.

**Figure 5:**
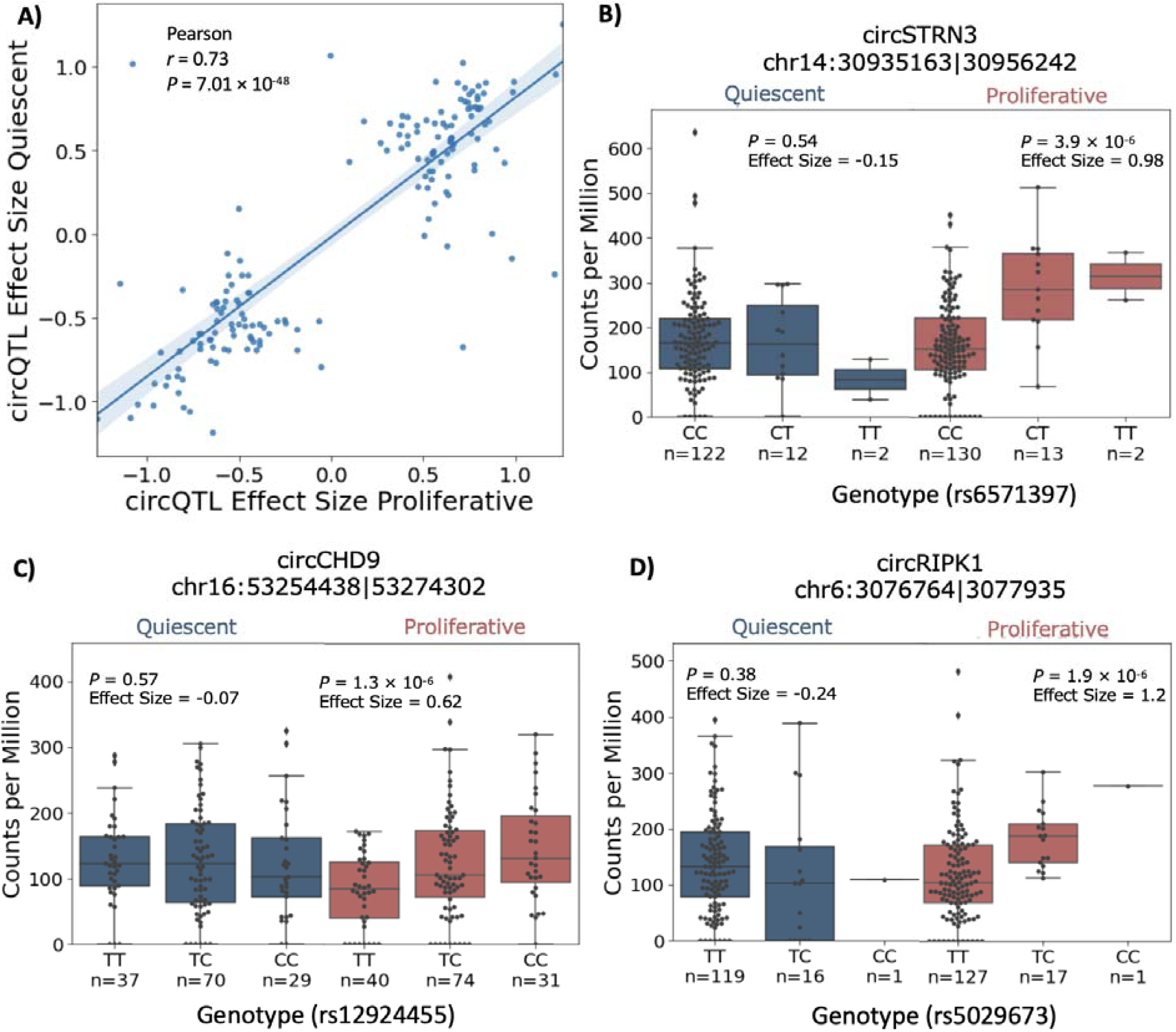
Phenotype-specific circQTL in SMCs. **A)** Pearson correlation between circQTL effect sizes in the proliferative and quiescent SMCs. **B-D)** Phenotype-specific circQTLs that had significant association in one phenotype but not the other.

### CircRNA isoform-specific circQTLs

We detected circRNA isoforms (circRNAs which differed in backsplicing sites but were derived from the same parent gene) arising from 30% of the parent genes (**Figure 2C**). Across both SMC phenotypes, we identified 34 parent genes which had two significant circRNA isoforms with QTLs. We identified no parent gene with more than two circRNA isoforms with QTLs. Therefore, we tested if an eSNP had opposite effect sizes on the expression of more than one circRNA isoform within a parent gene. In quiescent SMCs, we identified four eSNPs with circRNA isoform-specific direction. In proliferative SMCs, we identified two eSNPs with circRNA isoform specific direction (**Supplementary Table 4**). An example of an eSNP with circRNA isoform-specific direction is the SNP rs16875285. The C allele of this SNP is associated with lower expression of one circRNA isoform but higher expression of another circRNA isoform arising from the long non-coding RNA *GARS-DT* (**Figure 6A**). We obtained a similar result for circRNA isoforms in *TASOR2* (**Figure 6B**).

**Figure 6:**
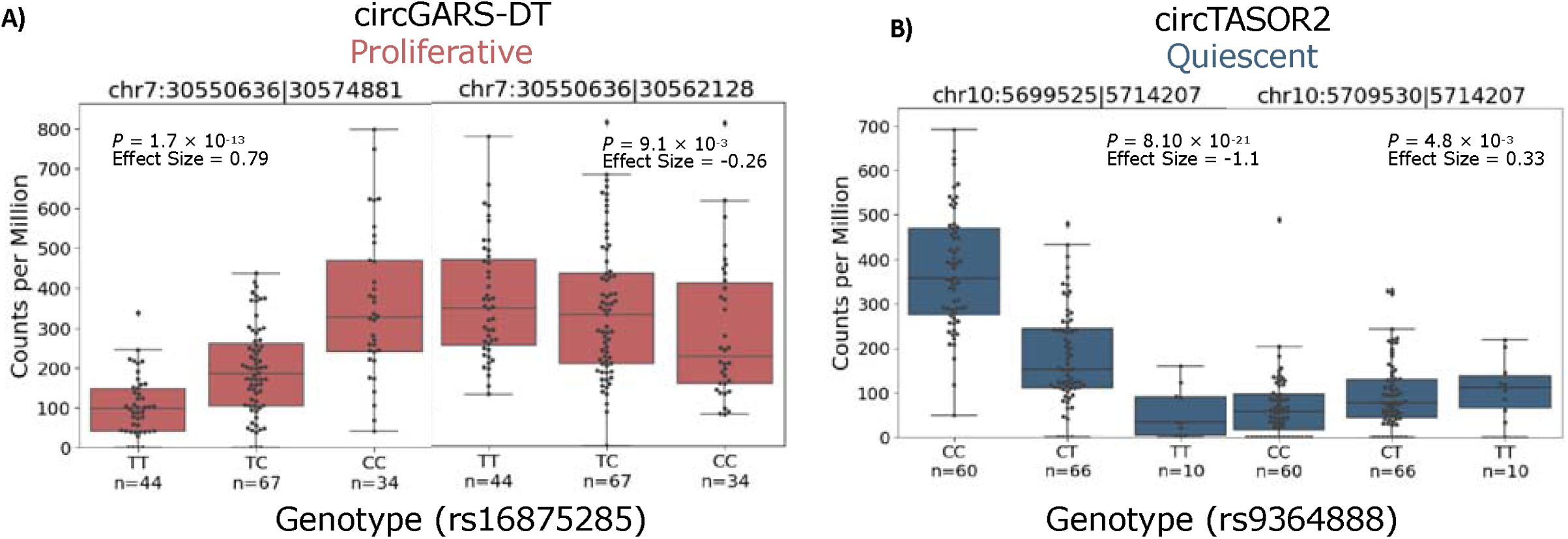
Genetic regulation of circRNA isoform abundance. **A**,**B)** Two examples where a genetic variant decreases the expression of one circRNA isoform but increases the expression of another circRNA of the same parent gene.

### CircQTL overlap with vascular disease loci

To uncover circRNAs that may be playing a role in the pathogenesis of diseases that involve SMCs, we tested if vascular disease related GWAS loci were associated with circRNA expression using a colocalization approach implemented in COLOC^18^ and eCAVIAR^19^. We tested for colocalization with CAD, myocardial infarction, stroke, aortic aneurysm, and blood pressure GWAS loci **(Supplementary Table 5)**. Across quiescent and proliferative SMCs, we identified nine circQTLs (*circZKSCAN1, circFOXK2, circANKRD36, circLARP4, circCEP85L, circGTF3C2, circPDS5A, circSLC4A7, and chr17:42610108*|*42659552*) which colocalized with CAD, myocardial infarction, and blood pressure GWAS loci (**Figure 7A, Supplementary Figure 3, Supplementary Table 6**,**7**). Three circQTLs were shared across multiple GWAS. We highlight the colocalization of *circSLC4A7* circQTL with high blood pressure GWAS loci (**Figure 7B**). Previously known, *SLC4A7* is a Na+, HCO_3_^−^ cotransporter that inhibits NO-mediated vasorelaxation, smooth muscle Ca^2+^ sensitivity, and higher blood pressure in mice.^20^ We demonstrated that the *circSLC4A7* eSNP shows significant association with circular but not mRNA expression (**Figure 7C**).

**Figure 7:**
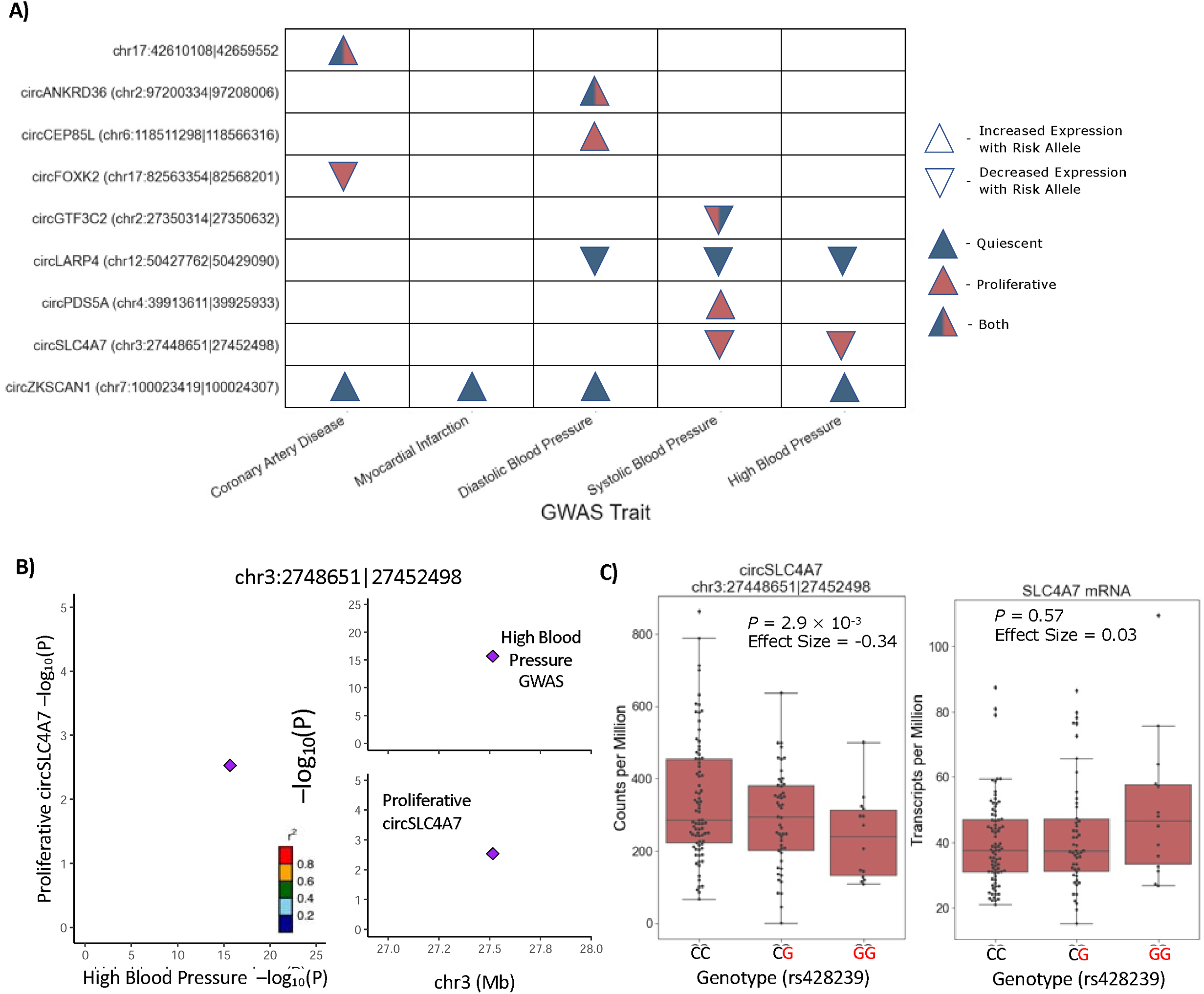
Cardiovascular GWAS loci associated with circRNA expression. **A)** Heatmap showing circQTL colocalized with cardiovascular GWAS loci across proliferative and quiescent SMCs using eCAVIAR and COLOC. **B)** circSLC4A7 circQTL signal colocalizes with the high blood pressure GWAS. The colocalized SNP, rs428239, is highlighted with a diamond. **C)** The risk allele, G, of rs428239 is associated with lower expression of circSLC4A7 but is not associated with mRNA SLC4A7 expression.

## DISCUSSION

Recent studies have identified roles for circRNAs in many complex diseases, including cardiovascular disease.^21^ It is often difficult to systematically measure and associate circRNA expression with a disease phenotype in a case-control setting. To overcome this, we used a systems genetics approach that utilizes naturally occurring randomly assigned genetic variants as putative causal mechanisms for changes in circRNA expression and disease phenotype. Previous GWAS have associated hundreds of loci with vascular-related disease traits where SMCs play key roles.^13,22^ The effect of vascular disease-related loci on circRNA expression has yet to be annotated. Thus, our study identified and characterized the effect of *cis*-acting genetic variation on circRNA expression in quiescent and proliferative aortic SMCs.

Using a unique source of 151 aortic SMCs from multiethnic donors, we identified 1,589 circRNAs of which 173 were differentially expressed between quiescent and proliferative SMCs. Although previous studies have performed differential expression of circRNAs between PDGF-BB treated and control vascular SMCs^11,12,23,24^, our study tested differences in circular RNA expression across SMCs cultured in the presence or absence of fetal bovine serum. While we did not treat our RNA with linear RNA degradation (Rnase R) enzyme like these other studies, we included many more samples (150 donors in two conditions). Our dataset provides unique insight into highly expressed circRNAs present across many individuals.

We identified 96 circRNAs associated with genetic loci which we term a circQTL. To our knowledge, this is the first study to perform genome wide circQTL analysis in SMCs; however, other studies have performed circQTL analyses in lymphoblastoid cell lines^25^ and dorsolateral prefrontal cortex samples.^26^ One previous study had identified that *circANRIL* had a circQTL in atherosclerotic plaques.^8^ Due to stringent expression threshold cutoffs, *circANRIL* was removed from our analysis. This implies that circQTL detected in atherosclerotic plaques could be due to expression in cells other than SMCs, cultured SMCs have a different expression profile for *circANRIL* than *in vivo* SMCs, or SMC phenotypic state that expressed *circANRIL in vivo* was not represented in our cultures.

Parent genes of the differentially expressed circRNAs were enriched in processes related to blood pressure: regulation of sodium ion transmembrane transport, regulation of heart contraction, and actin cytoskeleton organization. The circQTL parent genes were enriched for processes related to contraction: Rho protein signal transduction, sodium ion transmembrane transporter activity, and cAMP-mediated signaling. Overall, these results suggest that circRNAs play an important role in SMC processes and vascular disease traits. Previous studies have also demonstrated that circRNAs are relevant to SMC biology. For example, they showed that *circACTA2*^9^ and *circSFMBT2*^11^ may reduce SMC contraction. Although we detected *circACTA2* and *circSFMBT2*, these circRNAs were removed after stringent expression threshold cutoffs. Previous studies demonstrated that overexpression of *circMAP3K5*^12^ and *circLRP6*^10^ inhibited the proliferation of SMCs and could play a role in vascular pathogenesis. In our study, we did not find *circMAP3K5* and *circLRP6* differentially expressed between quiescent and proliferative cells.

To further characterize genetic architecture of circRNA expression in SMCs, we identified phenotype specific genetic regulation of *circSTRN3, circCHD9*, and *circRIPK1* expression where the effect size of eSNPs associated with circRNA expression differed between proliferative and quiescent cells. This is analogous to our previous studies which identified condition-specific effects of genetic regulation on mRNA expression in SMCs.^14^ Although further studies will be needed, phenotype specific genetic regulation of circRNA could be explained by transcription factor presence in one condition but not the other. Previous studies have demonstrated DNA-binding motifs targeted by transcription factors which, when bound selectively increase expression of circRNAs but not the host gene.^27^ In addition, we identified 6 SNPs which had distinct association directions with circRNA isoforms from the same gene. This could be explained by previous studies which found that competition of RNA pairing of inverted sequences surrounding a backsplicing sites could affect circRNA splicing selection.^28^

We also compared the genetic regulation of circRNA with the genetic regulation of mRNA. Consistent with other studies,^29^ we found a low correlation between circRNA expression and mRNA expression. Furthermore, we found that a circQTL locus is usually not associated with mRNA expression of the parent gene. Two previous circQTL studies had also showed that the effect direction of a SNP on a circRNA expression is mostly concordant with the effect size of the same SNP on the parent gene expression.^25,26^ Overall, these results suggest that circRNAs and mRNAs have distinct methods of expression regulation. Previous studies have suggested that SNPs associated with circRNAs may contribute to mechanisms associated with inverted sequences around a backsplicing site or canonical splicing sites.^26^

Lastly, we utilized a systems genetics approach to annotate the disease relevance of SMC circRNAs. We treated circRNA as an intermediate molecular phenotype bridging DNA variation with disease traits. Our previous study identified that 58 and 127 CAD loci were explained by changes in gene expression and alternative splicing, respectively. We demonstrated that circRNAs contribute to the genetic architecture of complex traits by colocalizing circQTL loci with vascular related GWAS loci. We identified nine circQTL loci – *circZKSCAN1, circFOXK2, circANKRD36, circLARP4, circCEP85L, circGTF3C2, circPDS5A, circSLC4A7*, and *chr17:42610108*|*42659552* – which overlap with CAD, myocardial infarction, and blood pressure GWAS loci. To date, no functional study of these circRNA transcripts have been carried out in the context of SMCs; however, current functional studies in cancer lines suggest that these circRNAs play a role in proliferation. Silencing of *circZKSCAN1* was shown to promote proliferation, migration, and invasion of hepatocellular carcinoma cell lines.^30^ *CircFOXK2* was shown to promote growth and metastasis of pancreatic ductal adenocarcinoma.^31^ Knockdown of *circANKRD36* promotes the apoptosis and inflammation of chondrocytes.^32^ *CircLARP4* inhibits cell proliferation and invasion of gastric cancer.^33^ Although no functional study has been reported for *circSLC4A7, SLC4A7* is a Na+, HCO_3_^−^ cotransporter that inhibits NO-mediated vasorelaxation, SMC Ca^2+^ sensitivity, and higher blood pressure in mice.^20^ To our knowledge, we are the first to characterize the *chr17:42610108*|*42659552* backsplicing site which is intergenic between the *TUBG* and *RETREG3* genes.

Overall, our results suggested that SMC circRNAs can play a role in the genetic risk for higher blood pressure, myocardial infarction, and CAD. Notably, for all circQTLs colocalized with vascular disease-related GWAS loci, we identified genetic regulation specific to circRNA expression but not its parent gene expression. This suggests that circRNAs explain an additional part of the genetic architecture of disease that is missed when looking at just mRNA gene expression.

## METHODS

### Cell culture, Genotyping, and RNA sequencing

As previously described, we collected SMCs from 151 multi-ethnic heart transplant donors. We cultured each of the 151 SMC samples in media with fetal bovine serum to represent proliferative SMCs and media without fetal bovine serum to represent quiescent SMCs.^14^ We performed RNA-seq of the ribosomal RNA-depleted total RNA. We prepared sequencing libraries with the Illumina TruSeq Stranded mRNA Library Prep Kit and Psomogen sequencing facility sequenced each sample to ∼100 million read depth with 150 bp paired-end reads. We genotyped each donor for ∼6.3 million SNPs with minor allele frequency >5%.^34^

### CircRNA Detection

We trimmed paired-end RNA-seq reads with low average Phred scores (<20) using Trim Galore and performed circRNA mapping and quantification with BWA and CircRNA Identifier 2 (CIRI2) in accordance with CIRI’s recommended pipeline with all default parameters except for -0, which outputs all backsplicing sites without filtering.^35,15^ We used the hg38 genome and GENCODE v32 annotations. CIRI counted circular junction counts, which is the number of reads overlapping a backsplicing site. We used circular junction counts as the quantitative measure of circRNA expression. CIRI also calculated the circular junction ratio, which is the ratio of circRNA expression to linear transcript expression as is given by:

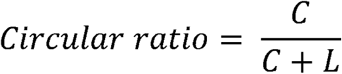

where C represents the number of circular counts and L represents the number of reads supporting a linear transcript at the junction.

### CircRNA Preprocessing

We considered a circRNA expressed if the junction count was ≥3 and the circular ratio was ≥0.05 in at least 20% of the samples of a particular phenotype. We used a threshold of counts ≥3 to reduce the number of circRNA false positives, and we used a threshold of ≥0.05 to reduce the possibility of mRNAs being miscounted as circRNAs. For downstream differential circRNA expression and circRNA quantitative trait loci mapping, we tested only expressed circRNAs. We classified circRNAs as exonic, intronic, and intergenic based on GENCODE v32 annotations. CIRI classified a circRNA as intronic or intergenic if at least one splice site was “intergenic” or “intronic” respectively; otherwise, CIRI classified a circRNA as exonic. CIRI determined a circRNA’s parent gene if the backsplicing site overlapped an intron or exon of an annotated gene. Intergenic circRNAs were not assigned a parent gene.

### Differential circRNA Expression

We performed differential circRNA expression between quiescent and proliferative SMCs using edgeR.^36,37^ In order to compare expression values across different samples, we applied trimmed mean of M values (TMM) normalization.^38^ Within edgeR, we used sex and the first four genotype principal components (PCs) as covariates. We considered a circRNA differentially expressed if the *q*-value false discovery rate was less than 0.05.^39^

### Circular RNA Quantitative Trait Loci Mapping

We tested the association of circRNA expression and ∼6.3 million variants. We term each significant association a circRNA quantitative trait locus (circQTL). To correct for library depth, we converted circRNA junction counts to counts per million (CPM) whereby a particular circRNA junction count was divided by the total junction counts in millions for that sample. We next normalized the CPM using TMM^38^ followed by inverse normalization. As covariates, we included technical confounders estimated through probabilistic estimation of expression residuals (PEER)^40^, 4 genotype principal components, and sex. From 2, 4, 6, 8, and 10 PEER covariates, we chose the number of PEER covariates that maximized the number of circQTLs. We performed circQTL detection using tensorQTL which implements a permutation scheme that accounts for linkage disequilibrium by approximating the distribution of *p*-values within a locus and subsequently accounts for genome wide testing using *q*-value correction.^15^ We considered a circQTL significant if the *q*-value false discovery rate was less than 0.05 We identified secondary circQTLs by rerunning permutation analysis but conditioning on the lead SNP of the locus.^17^ We tested variants within 500kb of either end of the backsplicing site. We term eSNP as the lead or most significant circQTL SNP in the locus.

### Phenotype-specific circQTLs

To determine phenotype-specific circQTLs, we tested if the eSNP for every significant circQTL had different effect sizes between proliferative and quiescent SMCs using a Z-score formulated as:

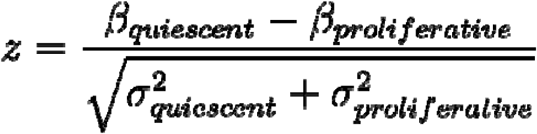

where *β* is the effect size of the eSNP and *σ* is the variance of the effect size.^41^ We determined a phenotype-specific circQTL if the Bonferroni-corrected P-value of the Z-test was < 0.05.

### CircRNA Isoform-specific circQTLs

We define circRNA isoforms when distinct backsplicing sites arise from the same parent gene. To identify if any eSNPs were associated with the expression of more than one circRNA isoform at a time, we tested for differences in effect size of an eSNP on two circRNAs deriving from the same parent gene. We performed analyses separately in each SMC phenotype. For every parent gene that had two significant circQTLs on different circRNA isoforms, we performed two statistical tests. First, we tested if the first eSNP had opposite effect size on the expression of the first circRNA isoform than the second circRNA isoform. Second, we tested if the effect of the second eSNP had opposite effect size on the expression of the first circRNA isoform than the second circRNA isoform. For each test we used the Z-score formulated as:

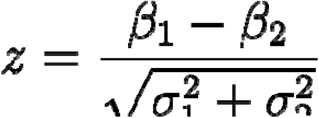

where *β*_*1*_ and *σ*_*1*_ are the effect size and variance of the primary circQTL and *β*_*2*_ and *σ*_*2*_ are the effect size and variance of the eSNP associated with another circRNA from the same parent gene. To account for the two tests performed per parent gene, we applied an initial Bonferroni correction by multiplying the *p*-value of the Z-test by two. Lastly, to account for multiple testing hypothesis across the genome, we applied a second Bonferroni correction to all parent genes tested.

To determine if there was a difference in directionality of effect sizes between an eSNP and two circRNAs, we took 90% confidence intervals of the effect size (β) of the eSNP on a circRNA. We classified an effect positive if the confidence interval did not include 0 and the Z-score was positive. Likewise, we classified an effect negative if the confidence interval did not include 0 and the Z-score was negative. Using the effect size annotations for directionality, we identified circRNA isoform-specific direction if the effect sizes differed in sign.

### Colocalization

To test whether the genetic variants were associated with circRNA expression but not mRNA expression, we performed coloclization of circQTL loci with eQTL loci using COLOC.^18^ From our previous study, we performed eQTL analysis using the same SMC samples and used these results for analysis.^14^ We performed colocalization with each significant circQTL and considered a locus colocalized if the posterior probability of colocalization (PPH4) was greater than 0.5.

To identify if circRNAs contribute to the genetic risk for vascular related diseases, we performed colocalization of circQTL loci with vascular disease-related GWAS loci using eCAVIAR and COLOC.^19,18^ We downloaded summary statistics for CAD, myocardial infarction, stroke, aortic aneurysm, and blood pressure GWAS catalog and UKBB **(Supplementary Table 5)**.^13^ For each circQTL, we considered variants within 500 kb of the eSNP, and at least one of the variants in the 500 kb window needed at least 10^−4^ GWAS *p*-value nominal significance. For each GWAS, we used a 1000G population matched linkage disequilibrium. For eCAVIAR, we set the maximum number of causal variants to 2 and considered variants colocalized if the colocalization posterior probability (CLPP) was greater than 0.01. For COLOC which uses a single causal variant assumption, variants were considered colocalized if the posterior probability of colocalization (PPH4) was greater than 0.5.

## Data Availability

RNAseq data is available at GEO with the accession number GSE193817.

