## Supplementary Figures for "Genetic Regulation of Circular RNA Expression in Human Aortic Smooth Muscle Cells and Vascular Traits"

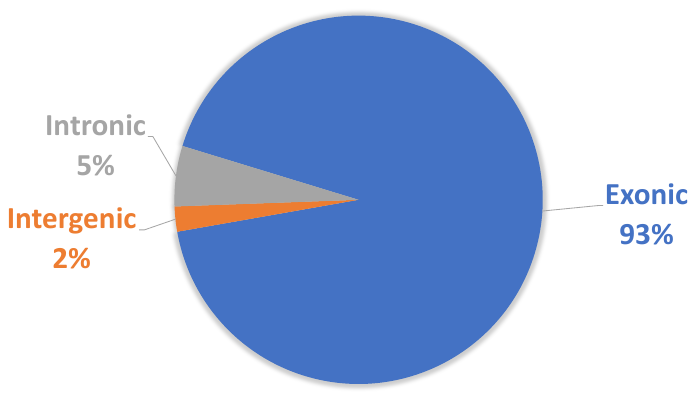

**Supplementary Figure 1: Classification of backsplicing sites.** We classified a circRNA intronic or intergenic if at least one splice site was “intergenic” or “intronic" respectively; otherwise, we classified a circRNA as exonic.

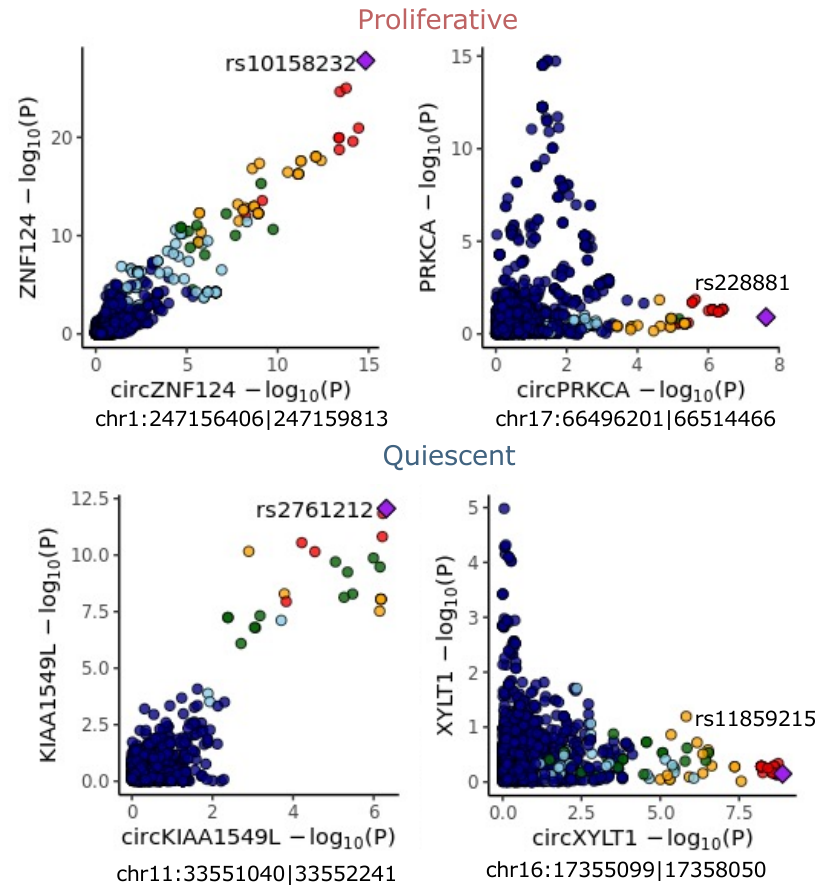

**Supplementary Figure 2: CircQTL loci association with parent gene mRNA expression.** The figures on the left are from circQTL and eQTL locus that passed 0.9 posterior probability of colocalization threshold where the lead SNP of the circQTL matches the lead SNP of the eQTL. The figures on the right have < 0.05 posterior probability of colocalization indicating different circQTL and parent gene eQTL SNPs.

**A)**

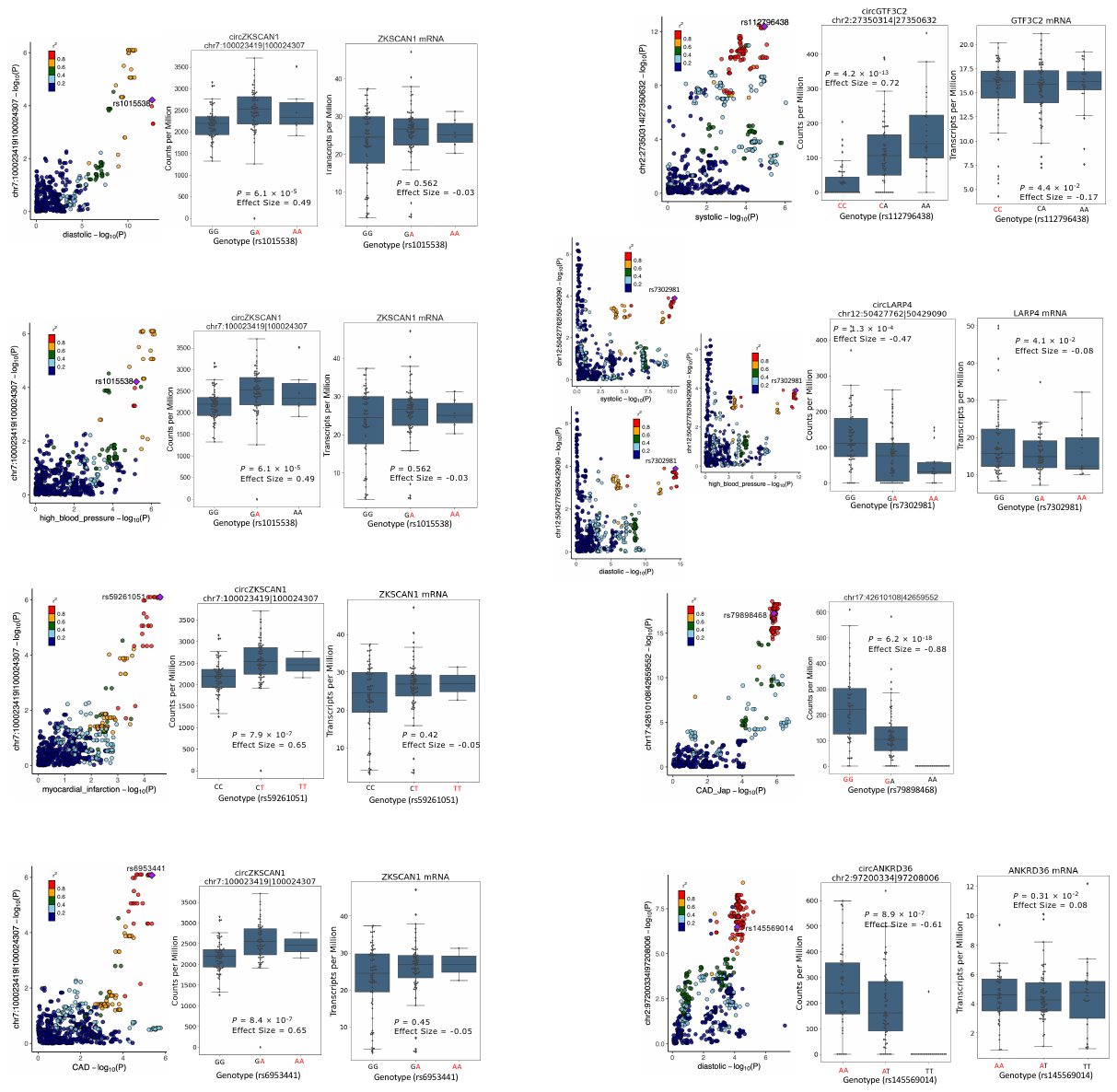

**B)**
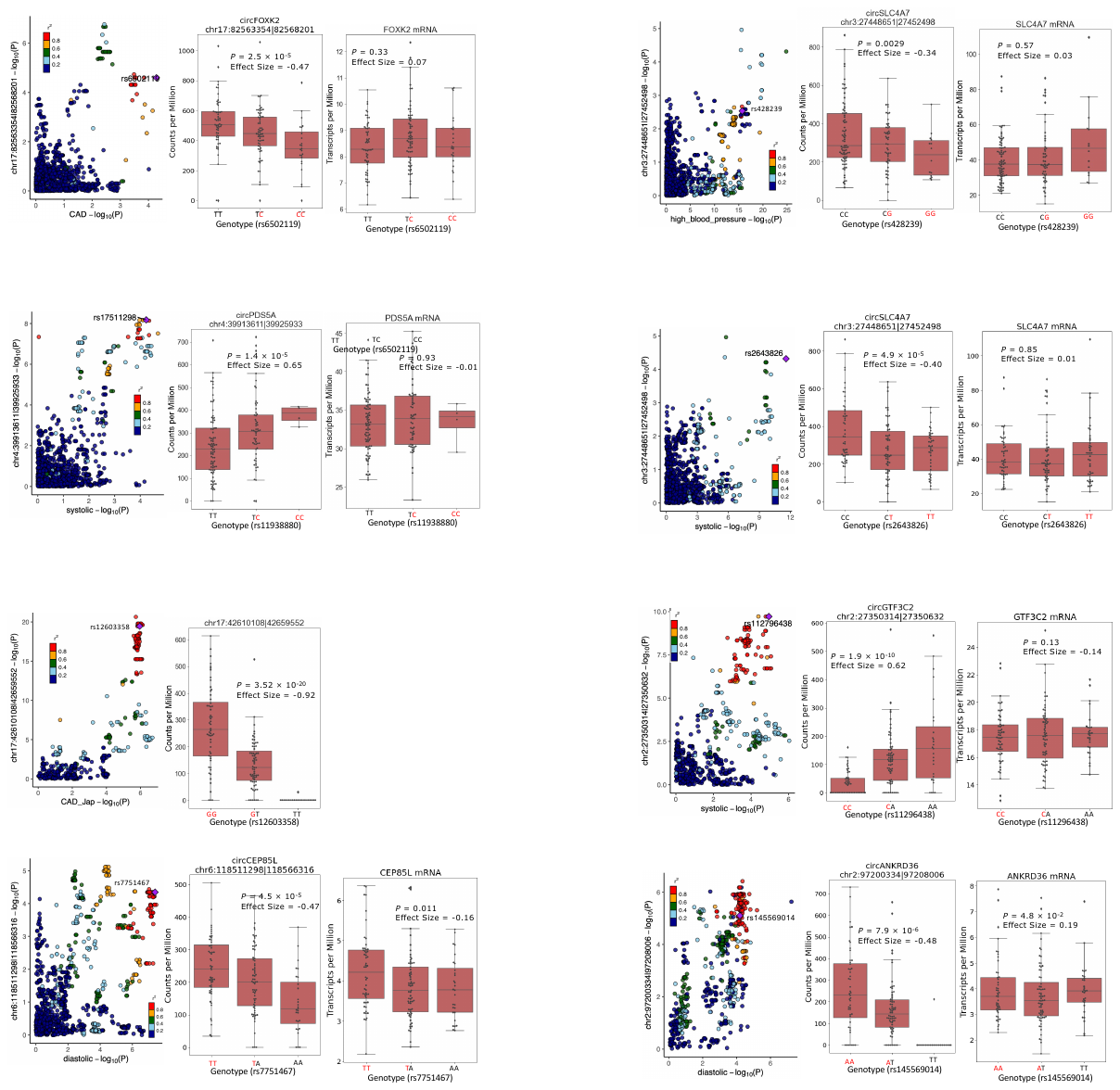

**Supplementary Figure 3: Additional cardiovascular GWAS loci associated with circRNA expression.** **A)** circQTLs colocalized in the quiescent SMCs are shown. For each row, the first figure represents the association between circQTL association against GWAS association. The colocalized SNP is indicated by a purple diamond. The second figure demonstrates the colocalized SNPs association with circRNA expression. The last figure represents the association of the same SNP with parent gene mRNA expression. The risk allele is highlighted in red. **B)** circQTLs colocalized in proliferative SMCs are shown.
